## Supplementary for "The H163A Mutation Unravels an Oxidized Conformation of the SARS-CoV-2 Main Protease and Opens a New Avenue for Anti-Viral Therapeutic Design"

### SUPPLEMENTARY INFORMATION

Norman Tran[1†], Sathish Dasari[2†], Sarah Barwell[1], Matthew J. McLeod[3], Subha Kalyaanamoorthy[2], Todd Holyoak[1*], & Aravindhan Ganesan[4*]

[1]Department of Biology, Faculty of Science, University of Waterloo, 200 University Avenue West, Ontario N2L 3G1, Canada

[2]Department of Chemistry, Faculty of Science, University of Waterloo, 200 University Avenue West, Ontario N2L 3G1, Canada

[3]Physics Department, Cornell University, Ithaca, NY 14853, USA

[4]ArGan's Lab, School of Pharmacy, Faculty of Science, University of Waterloo, 10A Victoria Street South, Kitchener, Ontario N2G 1C5, Canada

[*]corresponding authors

Aravindhan Ganesan,
Todd Holyoak,

[†]co-first authors

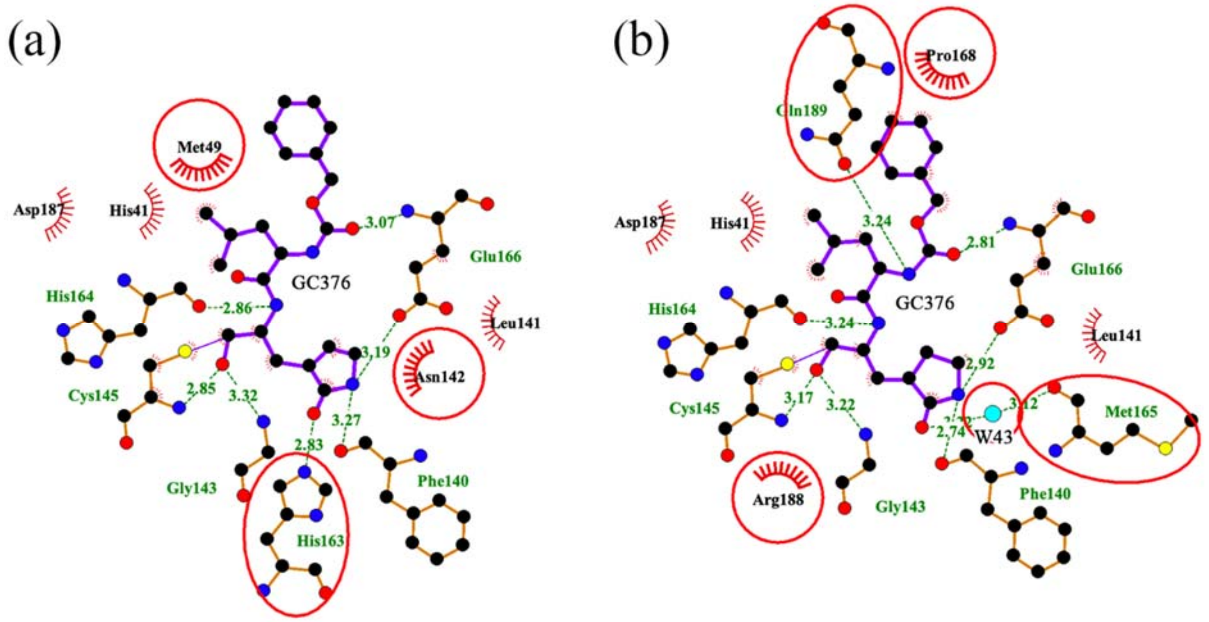

**Supplementary Figure 1 – 2D View of GC376-Mpro Interactions.** GC376 generally interacts with the same enzyme residues between the WT *(a)* and H163A mutant *(b)* enzyme. Many of these interactions are within the active-site pocket due to the nature of GC376 being a covalent inhibitor. GC376 also makes a hydrogen bond with H163 in the lateral pocket. This interaction is substituted with W43 in the H163A structure. Circled residues highlight differences between the two structures. This figure was made with LigPlot+.[68]

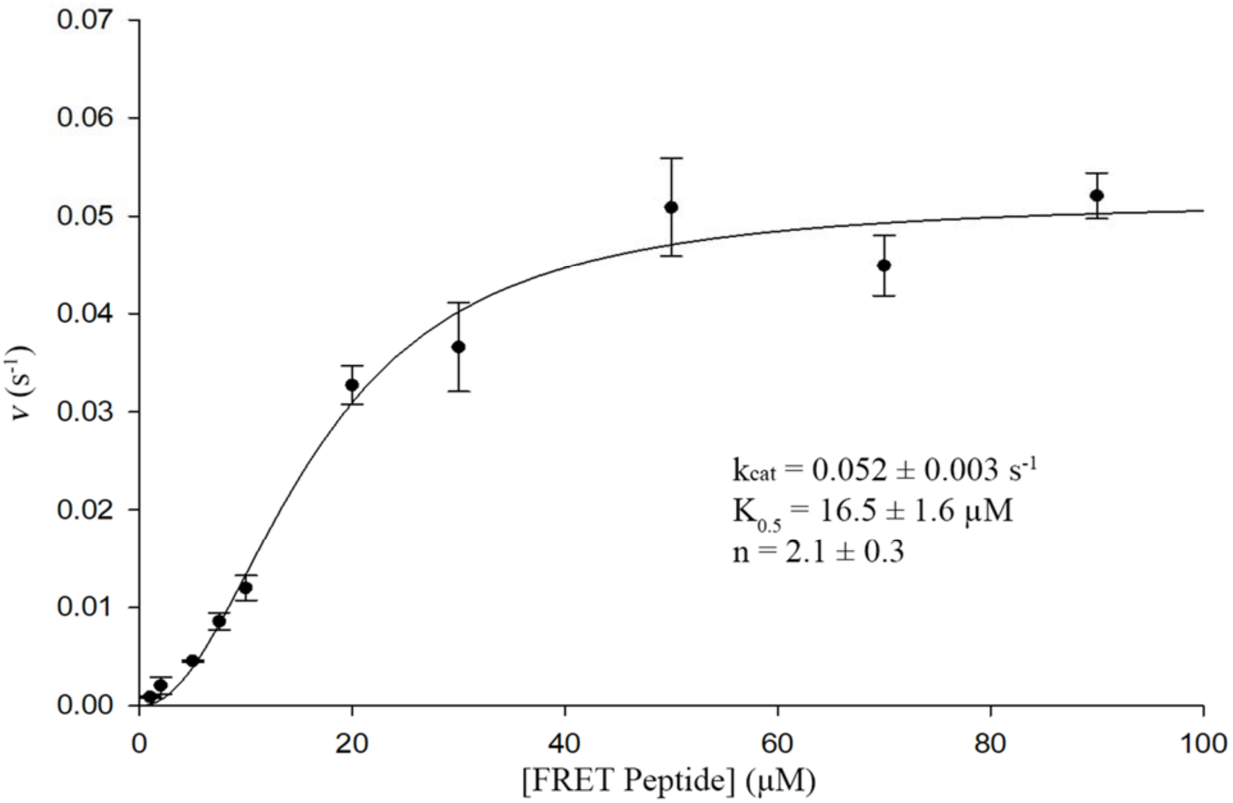

**Supplementary Figure 2 – Michaelis-Menten Curve for WT Mpro.** Initial steady-state rates for WT Mpro were measured via a fluorescent-based kinetic assay at various concentrations of fluorescent peptide (number of replicates = 3). These data fit well to the Hill equation and showed positive kinetic cooperativity between the active sites within each monomer. Error bars represent the standard deviation.

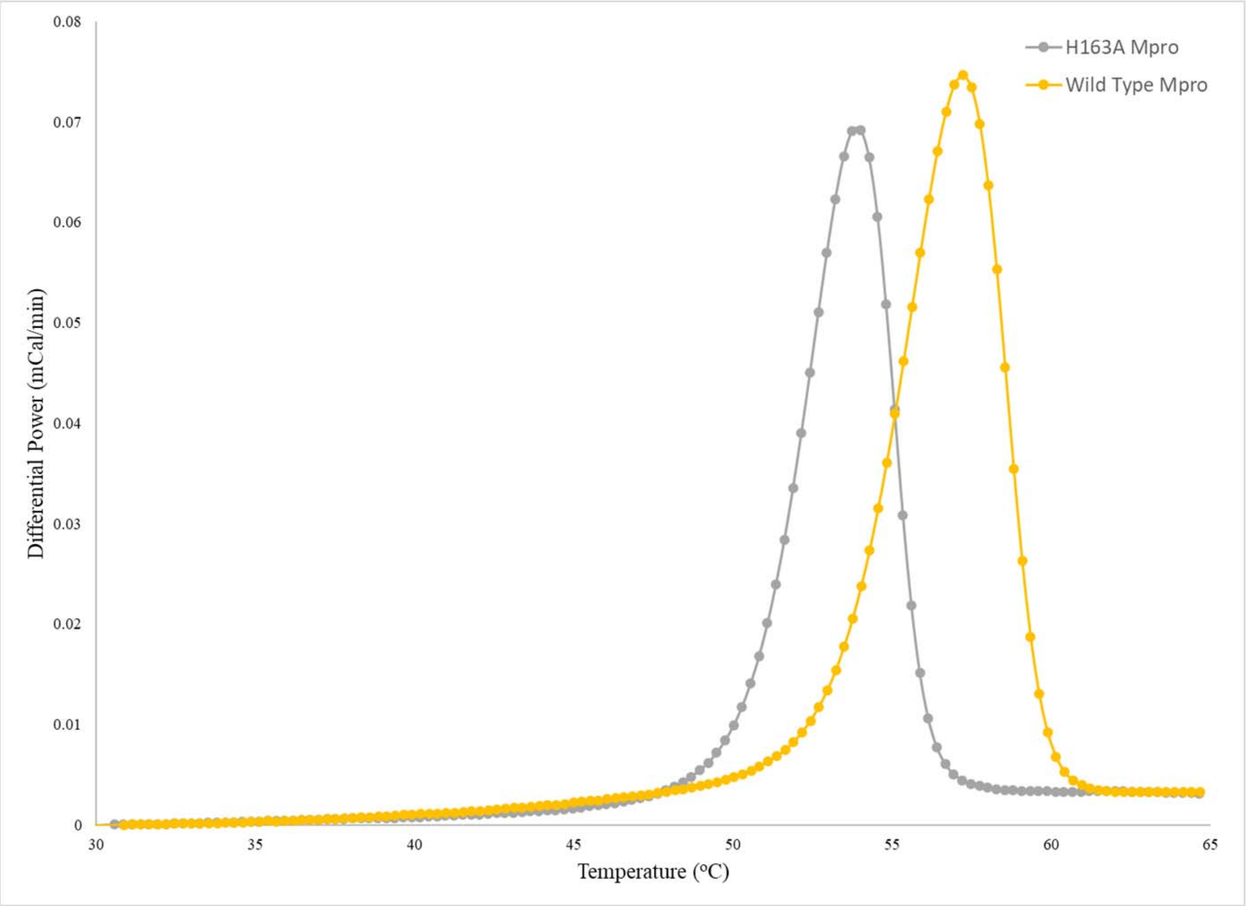

**Supplementary Figure 3 – Differential Scanning Calorimetry Thermograms of WT and H163A Mpro.** The thermograms of WT and H163A Mpro show significant differences in thermal stability. Despite the peak shape fitting well to a concerted, oligomeric unfolding model, thermodynamic and mechanistic information about the enzymes' unfolding cannot be extracted from these data.

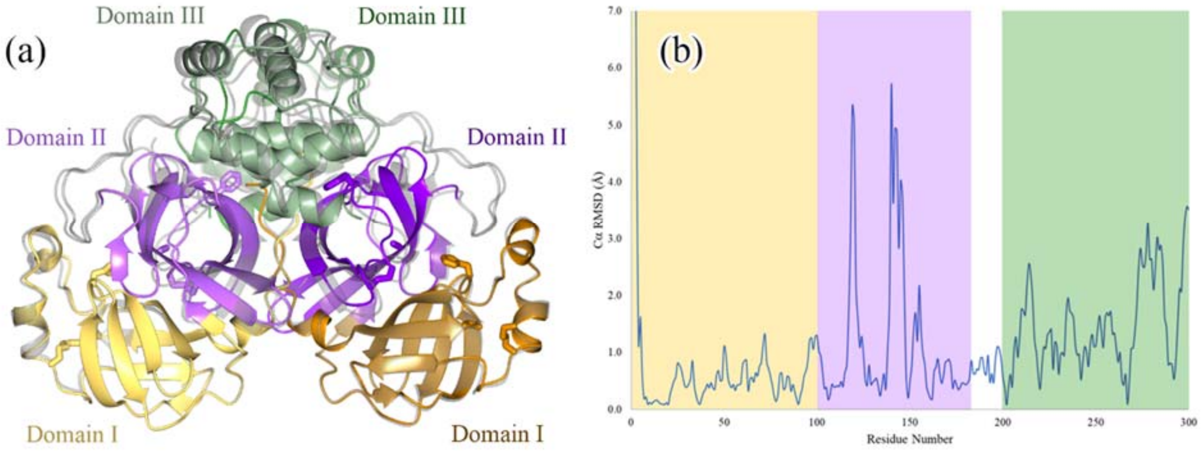

**Supplementary Figure 4 – Global Comparisons Between Apo WT and Apo H163A Mpro Structures.** *(a)* Global differences are seen when comparing the structures of the apo WT (grey) and apo H163A mutant. This is most noticeable globally in Domain III (pale green/forest green), where several of the helices in the H163A structure are displaced relative to their positions in Domains I and II of WT Mpro. *(b)* A comparison of Cα root-mean-squared deviation (RMSD) values between the WT and mutant residues. Cα RMSD values were calculated using Chimera.[54] Due to the structural asymmetry between the two molecules in the asymmetric unit, Cα RMSD values were only calculated for chain B as it showed more structural deviation compared to chain A. The figure is colored by the same scheme as in (a). The most notable spikes in Cα RMSD values for each domain from the N- to C-terminus correspond to a repositioning of the N-terminus in Domain I, rearrangement of the active-site and surrounding loops in Domain II, and an overall displacement of helices in Domain III.

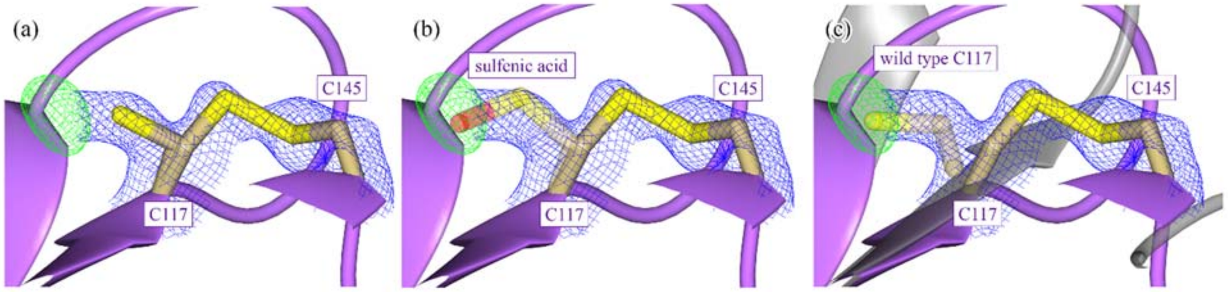

**Supplementary Figure 5 – Accounting for the Positive Difference Density at C117 in the Apo H163A Mutant Structure.** *(a)* There is strong positive $F_o$-$F_c$ density (4.0 σ $F_o$-$F_c$; 1.2 σ $2F_o$-$F_c$) adjacent to the sulfur atom of C117 when the disulfide bond between C117 and C145 is broken. It is unclear whether this positive density is due to an oxidized C117 as sulfenic acid *(b)* or if a minor proportion of the beta strand containing C117 relaxes into its WT conformation (grey; PDB 7BB2), placing the WT C117 into the positive density *(c)*. Because of this ambiguity and lack of direct evidence for either possibility, C117 is modelled as a reduced cysteine in the H163A mutant structure.

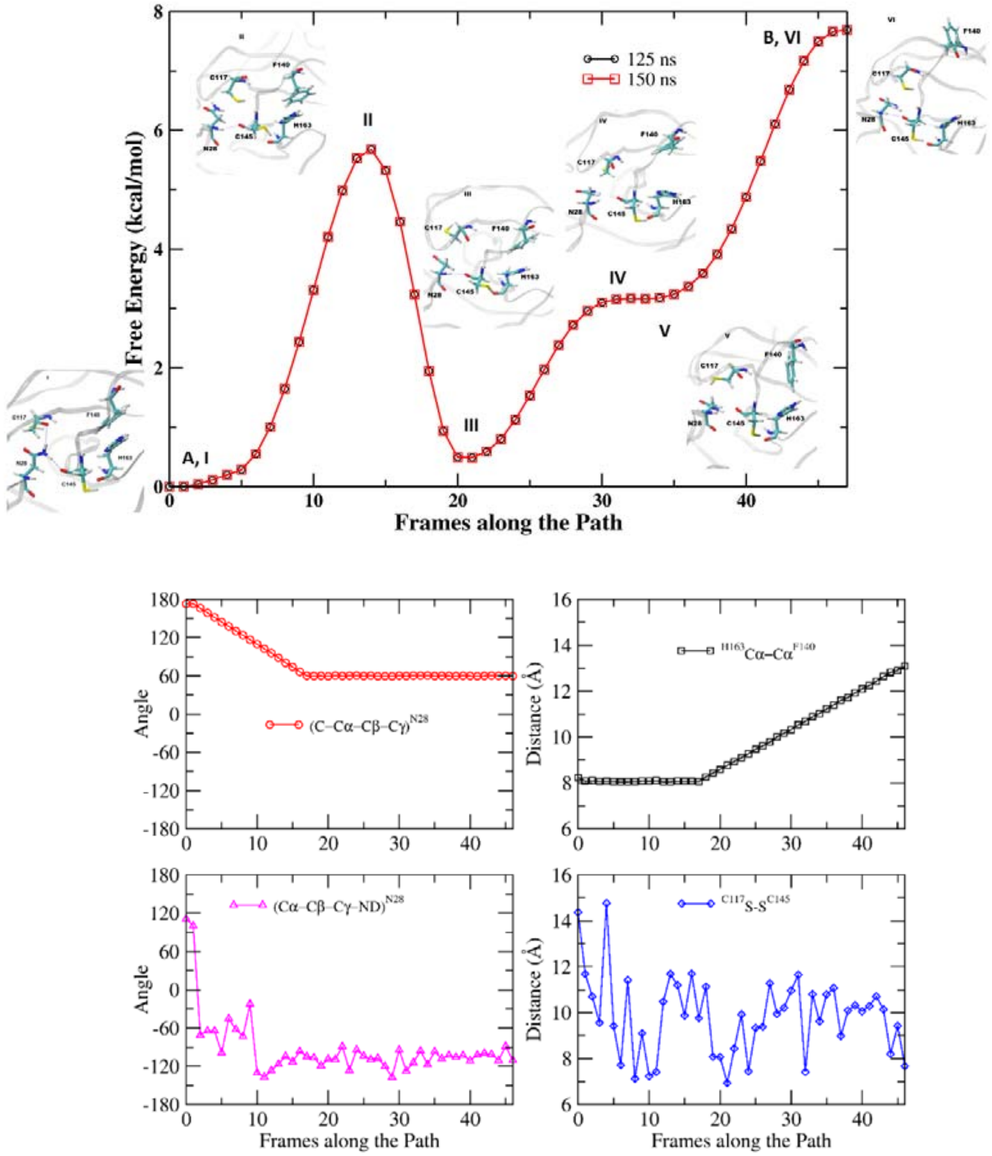

**Supplementary Figure 6 – MFEP for WT model**. *(top half)* (A, I) Conformation of WT structure; (II) transition state corresponding to the rotation of the N28 side chain; (III) minimum corresponding to the conformation after rotation of N28 dihedral; (IV) transition state corresponding to the dissociation of π-π stacking; (V) minimum corresponding to the C-H-π interaction between F140 and H163; (B,VI) Conformation corresponding to the mutated crystal structure. (*bottom half*) Change in dihedral angles of the N28 side chain happens first then Cα distance between H163 and F140 increases. The S-S distance between C117 and C145 goes from 14 to 8 Å along the MFEP.

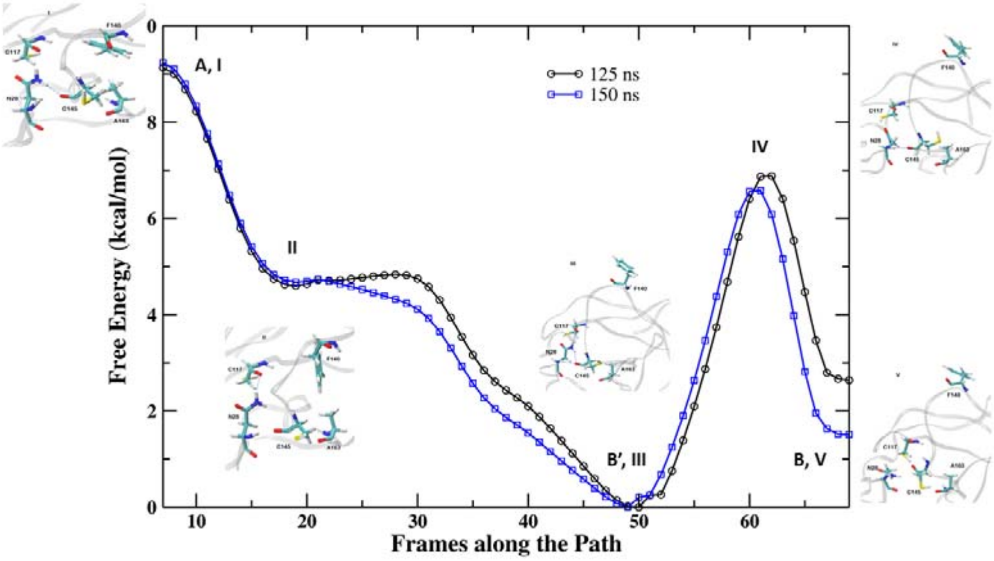

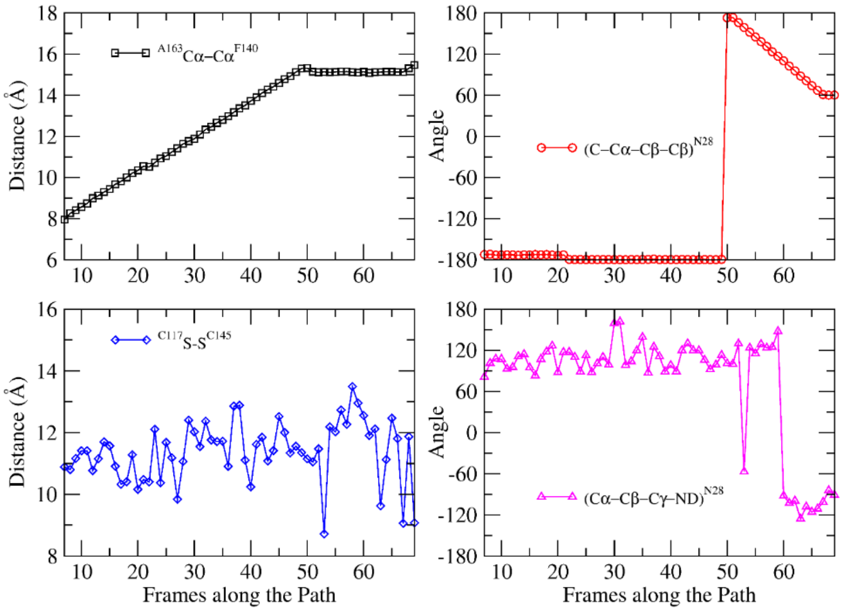

**Supplementary Figure 7 – MFEP for H163A model**. (*top half*) (A, I) Conformation corresponding to the WT structure for H163A mutated system; (II) Intermediated conformation along the dissociation of Cα distance between A163 and F140; (B', III) Conformation corresponding to elongated Cα distance between A163 and F140; (IV) Transition state conformation corresponding to the rotation of N28 side chain; (B, V) Conformation corresponding to the mutant crystal structure. (*bottom half*) Cα distance between H163 and F140 increases first followed by rotation of the N28 side chain. The S-S distance between C117 and C145 goes from 11 to 9 Å along the MFEP.

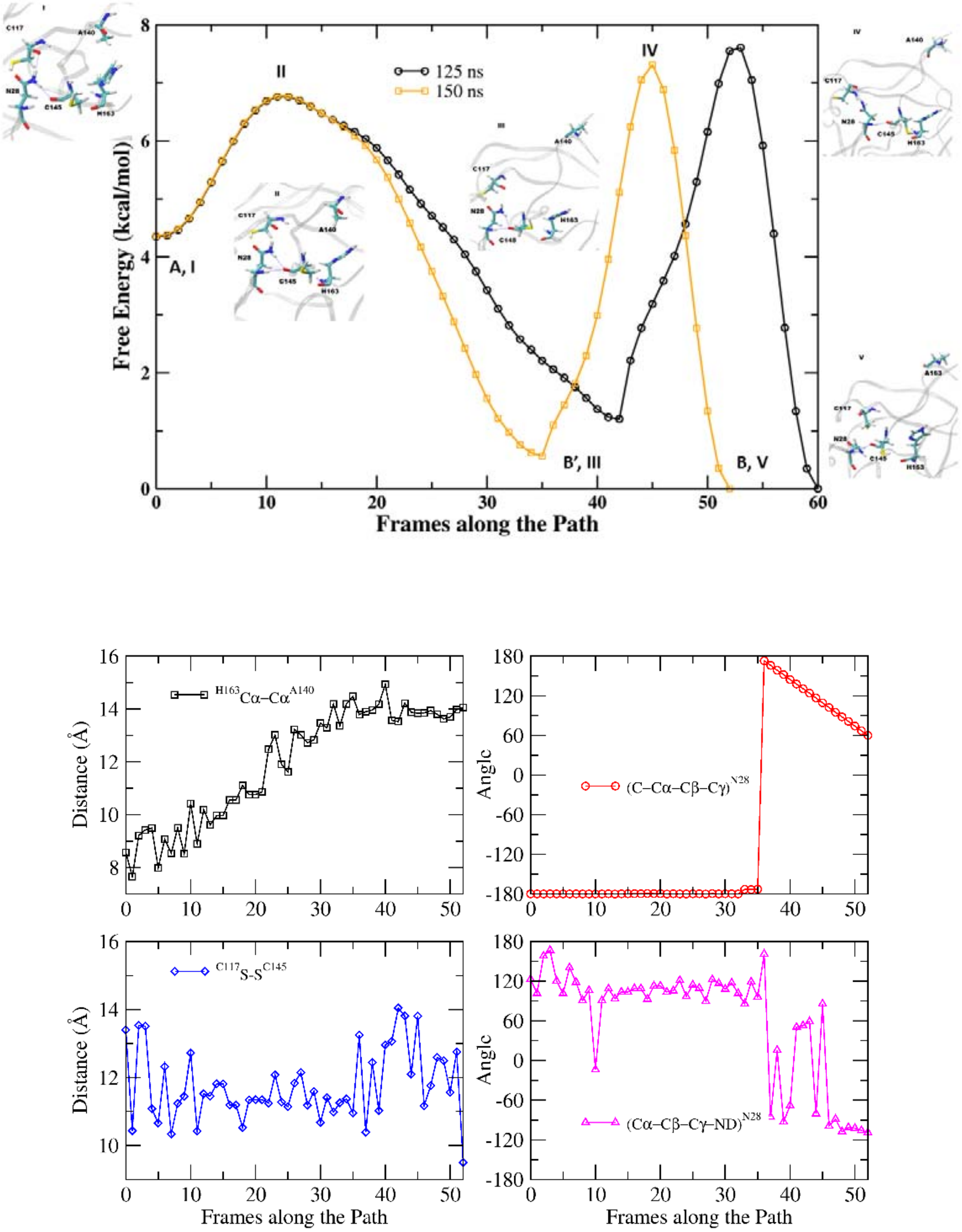

**Supplementary Figure 8 – MFEP for F140A model**. (*top half*) (A, I) Conformation corresponding to the WT structure for F140A system; (II) Transition state conformation along the dissociation of Cα distance between H163 and A140; (B', III) Conformation corresponding to elongated Cα distance between H163 and A140; (IV) Transition state conformation corresponding to the rotation of the N28 side chain; (B, V) Conformation corresponding to the mutant crystal structure. (*bottom half*) Cα distance between H163 and F140 increases first then the rotation of the N28 side chain occurs. The S-S distance between C117 and C145 goes from 13 to 9.5 Å along the MFEP.

.

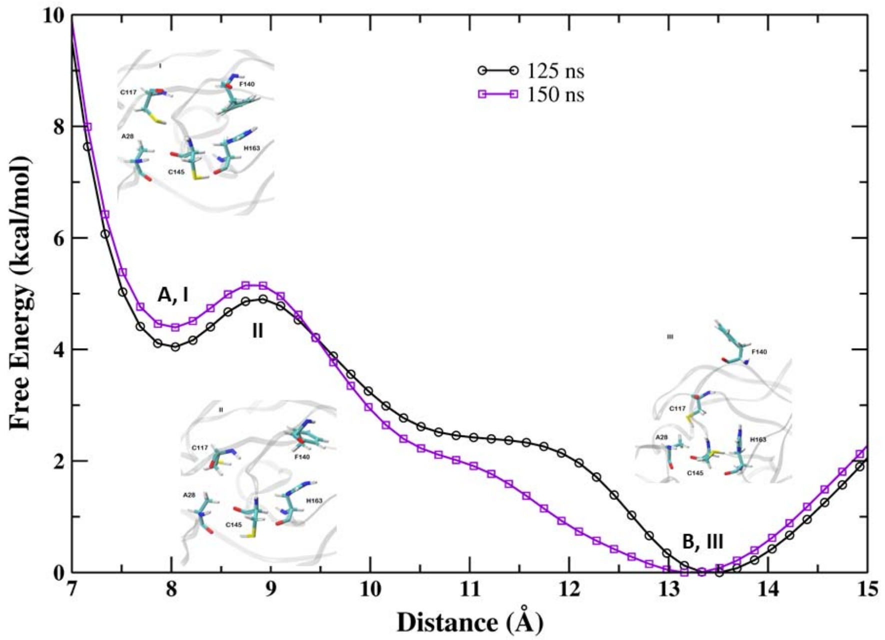

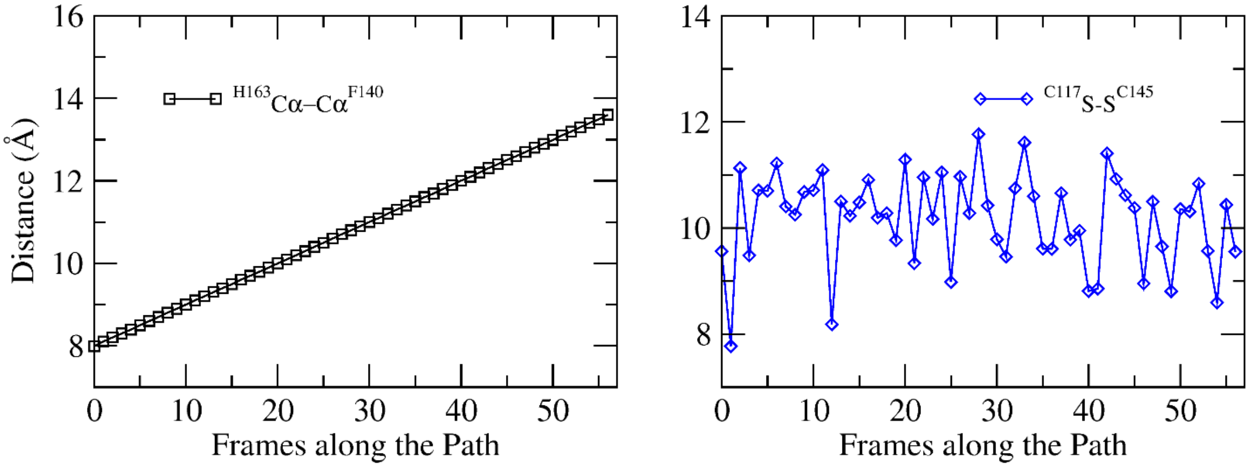

**Supplementary Figure 9 – MFEP for N28A model**. (*top half*) (A, I) Conformation corresponding to the WT structure for N28A system; (II) Transition state corresponding to the dissociation of π-π stacking and dissociation of Cα distance between H163 and F140; (B, III) Conformation corresponding to the mutant crystal structure. (*bottom half*) S-S distance between C117 and C145 fluctuates between 9 and 11 Å along the MFEP path.

**Supplementary Table 1 – Data Collection and Refinement Statistics.**

| | H163A Mpro with GC376 (**8DD6**) | Apo H163A Mpro (**8DDL**) |
|---|---|---|
| Wavelength (Å) | 1.1271 | 0.9686 |
| Resolution range | 54.97 – 2.30 (2.38 – 2.30) | 56.54 – 1.94 (2.01 – 1.94) |
| Space group | I2 | $P2_12_12_1$ |
| Unit cell | 44.92 52.84 111.74 β = 100.3° | 67.83 101.46 102.34 |
| Total reflections | 74279 (3750) | 711101 (36076) |
| Unique reflections | 11561 (558) | 52678 (2569) |
| Multiplicity | 6.4 (6.72) | 13.50 (14.04) |
| Completeness (%) | 99.76 (98.27) | 99.48 (95.00) |
| Mean I/σ(I) | 5.8 (0.5) | 6.4 (0.5) |
| Wilson B-factor | 36.06 | 28.34 |
| $R_{merge}$ | 0.175 (0.895) | 0.169 (1.082) |
| $R_{meas}$ | 0.191 (0.971) | 0.176 (1.122) |
| $R_{pim}$ | 0.075 (0.374) | 0.047 (0.297) |
| $CC_{1/2}$ | 0.991 (0.814) | 0.996 (0.759) |
| Number of reflections used in refinement | 11560 (1134) | 52608 (4940) |
| Number of reflections used for $R_{free}$ | 579 (46) | 2661 (241) |
| $R_{work}$ | 0.1973 (0.2767) | 0.1683 (0.2199) |
| $R_{free}$ | 0.2504 (0.3570) | 0.2067 (0.2679) |
| Number of atoms | 2499 | 5174 |
| Protein | 2416 | 4729 |
| Ligands | 4 | 69 |
| Water | 79 | 376 |
| B-factors (Å$^2$) | 43.51 | 35.35 |
| Protein | 43.57 | 34.76 |
| Ligands | 49.35 | 53.16 |
| Water | 41.34 | 40.90 |
| Root mean square deviations | | |
| Bonds (Å) | 0.002 | 0.005 |
| Angles (°) | 0.49 | 0.76 |
| Rotamer outliers (%) | 0 | 0.56 |
| Clashscore | 1.48 | 5.05 |
| Ramachandran (%) | | |
| Favored | 98.68 | 97.81 |
| Allowed | 0.99 | 2.19 |
| Outliers | 0.33 | 0 |

**Supplementary Discussion 1 – Summary of the Working Model for Generating the Oxidized, Disulfide-Bonded Mpro Conformation.**

In summary, the specifics of the working model for the increased energetic favorability of the oxidized structural state are as follows:

- The H163A mutation results in the loss of the face-to-face π-stacking interaction between H163 and F140 (*Fig. 5a*)
- The energetic barrier between the "in" and "out" conformations of F140 and the S139-S147 loop is reduced due to destabilization of the "in" conformation (*Fig. 6*)
- As the F140 side chain normally resides in a hydrophilic environment, upon adopting this "out" position, residues S147, H172, Y118, and Y126 rearrange to form a new hydrogen bonding network alongside two new water molecules (*Fig. 5c*)
- Rearrangement of the active-site loop (S139-S147) allows for the active-site C145 to come in close proximity to C117 (compare *Fig. 1b* with *Fig. 3a*)
- A disulfide bond between C117 and C145 forms to stabilize this "out" conformation and is structurally concomitant with the rotation of the N28 side chain (*Fig. 3a*)
- A NOS bridge between K61 and C22 is seen in one of the two protomers (*Fig. 4ab*)
- The N-terminus of one of the protomers is threaded 90$^{o}$ from its wild type orientation as the active-site loop, and in particular F140, sterically occludes the original position of the N-terminus (*Fig. 4cd*)
- The disulfide bond between C117-C145 is reversible under reducing conditions, as evident by kinetic data (*Supplementary Fig. 2*) and the mutant structure in complex with GC376 (*Fig. 2*)
- An energetically relaxed, wild-type-like conformation of the beta strand containing the active-site cysteine (S144-V148) is seen when the disulfide bond is broken (*Fig. 3bc*)
- The "in" to "out" conformation change rarely occurs the wild type enzyme as, in the presence of reducing agent (i.e. in normal physiological conditions), the C117-C145 disulfide bond is readily broken and the S139-S147 loop is restructured back into the "in" conformation as F140 is easily stabilized by H163 (*Fig. 6*)
- *In silico* metadynamics simulations of the F140A and N28A mutants showed similar conformational behavior to the H163A mutant (*Fig. 6*), suggesting that these residues play an important role in stabilizing the WT conformation
